# Opportunistic pathogens are prevalent across the culturable exogenous and endogenous microbiota of stable flies captured at a dairy facility

**DOI:** 10.1101/2024.11.04.621909

**Authors:** Andrew J. Sommer, Courtney L. Deblois, Andrew D. J. Tu, Garret Suen, Kerri L. Coon

**Author notes:** Corresponding author: Kerri L. Coon, 1550 Linden Drive, Madison, WI 53706.

## Abstract

Stable flies in the genus *Stomoxys* are highly abundant, blood-feeding pests on dairy farms; however, their role in the carriage and potential transmission of pathogens is largely understudied. Here, we report on the frequency and distribution of culturable bacteria collected from *Stomoxys* flies captured in free stall barns and nearby calf hutches over a three-month period on a focal research farm in Wisconsin, USA. Mastitis-associated bacterial taxa, including *Staphylococcus*, *Escherichia*, *Enterobacter*, and *Klebsiella* spp., were frequently isolated from pooled samples of the internal or external portions of the flies. Conversely, selective enrichment protocols from these samples yielded only a single isolate of *Salmonella* and no enterohemorrhagic *Escherichia coli* O157. Neither trap location nor time of capture had a significant impact on the observed frequency of most bacterial genera isolated from the flies. Our results confirm that *Stomoxys* flies harbor both mastitis-associated bacterial taxa and bacterial taxa associated with opportunistic infections in humans. Further research into the transmission of fly-associated microbes could be important in the control of mastitis or other bacterial diseases on dairy farms.

## Introduction

Biting stable flies (*Stomoxys* spp.) are major economic pests of dairy cattle and other livestock [1]. As obligate blood-feeders, male and female adult flies acquire daily nutritional bloodmeals via stabbing mouthparts, often causing serious physical irritation and stress to the host (Foil and Hogsette, 1994). Biting activity by flies is directly linked to decreased productivity, including a reduction in calf-weight gain and milk production in lactating dairy cows [1]. *Stomoxys* fly control is often very difficult, as high fly populations are easily sustained through the constant access of available hosts for bloodmeals, as well as access to preferred breeding sites such as manure and silage [2,3].

The close association of flies with cattle manure, a major reservoir of fecal-borne pathogens [4], suggests a role for flies in bacterial disease transmission. Adult flies are highly mobile and may mechanically disperse ingested bacteria through regurgitation during bloodmeals, defection, or via carriage on external surfaces [5–7]. Of particular concern is the potential role of *Stomoxys* flies in the dispersal of bovine mastitis and enteritis pathogens, which pose significant economic and health threats to both cattle and dairy workers. Cattle also serve as a major reservoir for bacterial pathogens responsible for gastrointestinal infections in humans including *Salmonella* and enterohemorrhagic *E. coli* (EHEC), which are transmitted to the environment via fecal shedding [8–10]. In addition, both *Salmonella* and ETEC are major causative agents of bovine neonatal enteritis and diarrhea – the leading causes of mortality for calves on dairy farms [9,11]. Environmental mastitis pathogens, including strains of *Enterobacteriaceae* (*Klebsiella*, *Escherichia*, *Enterobacter* spp., etc.), non-aureus staphylococci (NAS), and *Streptococcaceae* are widespread in manure and soiled bedding and can cause intramammary infections after contact by a susceptible dairy cow [12]. Infection by either *Salmonella* or mastitis pathogens can reduce both milk quality and total yield, with severe cases resulting in the culling of animals [13,14].

Although prior studies on the culturable microbiota of *Stomoxys* flies have identified the presence of *E. coli*, *Salmonella*, *Staphylococcus*, and other potential bacterial pathogens [15–17], population-wide incidence rates of bacterial carriage by *Stomoxys* flies over time and across different locations within dairy facilities are still largely unknown. As such, we have a limited understanding of the potential role of biting flies in disease transmission. Recently, we performed the first culture-independent characterization of bacterial communities in adult stable flies and cattle manure collected longitudinally across two dairy research farms in South Central Wisconsin, USA [18]. Many of the same bacterial strains (amplicon sequencing variants) were detected in both flies and manure samples, including taxa associated with mastitic cows housed in the same facilities (*Escherichia, Klebsiella*, *Staphylococcus* spp.). Mastitis associated taxa were found in significantly high abundances in flies, relative to manure, and viable colonies were readily isolated from fly samples [18]. However, while this study provides the first evidence to definitively support the potential of *Stomoxys* flies to transmit mastitis associated bacteria *in situ*, our sequencing and culturing methods were not designed to differentiate *Salmonella*, ETEC, and EHEC from other *Enterobacteriaceae* and *E. coli* strains [19]. Moreover, that study only considered flies collected directly from, or adjacent to, heifer housing structures and did not include those collected from nearby calf hutches, where enteric pathogens like *Salmonella* and ETEC may be more abundant in the environment.

The overall objectives of this study were to quantify and compare incidence rates of enteric and mastitis associated bacterial pathogens isolated from *Stomoxys* flies collected from a focal dairy facility in South Central Wisconsin. Flies were trapped from both heifer and calf housing structures over a three-month period during peak fly season. Culture-based and molecular methods were then used to screen fly-derived bacterial populations for taxa of interest. Our results provide the first comprehensive examination of clinically relevant bacterial taxa cultured from *Stomoxys* flies in a dairy environment and have important implications for our understanding of the role of these, and other flies, in shaping pathogen persistence and transmission in agricultural settings.

## Materials and Methods

### Field sampling and pooling design

Field work was performed at the Emmons Blaine Arlington Dairy Research Center, a free-stall dairy research facility located in Arlington, WI, USA. Flies were caught on adhesive alsynite fiberglass traps (Olsen Products Inc., Medina, OH), which selectively attract *Stomoxys* flies through reflection of ultraviolet light [20]. This included four traps placed around the perimeter of two main barn structures and a single trap placed in the center of an outdoor calf hutch area located ~60 m from the barns. Adhesive trap liners were retrieved and replaced weekly from July-September 2021. Retrieved liners were immediately bagged and stored at −20 °C until further processing in the laboratory.

Biting flies within the genus *Stomoxys*, which have pronounced piercing mouthparts, were identified with the assistance of taxonomic keys available in the Manual of Nearctic Diptera [21]. Ethanol-sterilized featherweight tweezers (DR Instruments Inc. DRENTF-II, Bridgeview, IL) were used to carefully remove flies from the adhesive liners. Flies retrieved from the four barn traps were then randomly sorted into eight early-season pools (July capture dates) and eight late-season pools (August and September capture dates), while flies retrieved from the calf hutch area trap were randomly sorted into four early-season and three late-season pools. All barn-derived fly pools consisted of 10 flies; calf hutch area derived fly pools consisted of 8-11 flies.

### Culturing and enrichment of fly-associated microbes

Fly pools were vortexed gently for 40 seconds in 10 ml sterile PBS-T (1X PBS + 0.01% Tween 80) followed by removal of the flies to generate an external fly-bacterial sample. These samples were then centrifuged (20 min; 3200 rcf) and the resulting pelleted cells were removed and resuspended in 1 ml sterile 1X PBS. The pooled flies from these samples were surface sterilized via successive washes in 70% ethanol, 0.05% bleach, and water, and homogenized by bead-beating with 3 x 5 mm stainless steel beads (Qiagen, Hilden, Germany) in 1 ml sterile 1X PBS to produce internal fly-bacterial suspensions.

For both internal and external fly-bacterial suspensions, 50 μl aliquots were directly plated as serial dilutions on Eosin-Methylene Blue (EMB) (BD, Franklin Lakes, NJ), MacConkey (Hardy Diagnostics, Santa Maria, CA), Mannitol Salt (Neogen, Lansing, MI), and trypticase soy blood agar (5% sheep blood) (Thermo Scientific, Waltham, MA) and incubated for 48 h at 37 °C. The remaining volume of each suspension was then grown overnight in Brain Heart Infusion (BHI) broth to non-specifically enrich bacteria from fly pools. Enrichment cultures were subsequently plated as serial dilutions on BHI agar (Dot Scientific, Burton, MI) and incubated for 48 hours at 37 °C.

### Bacterial 16S rRNA sequencing and analysis

Bacterial colonies displaying various phenotypes were selected from each agar plate to obtain a representative collection of the culturable fly microbiota. Colonies selected for further analysis were streaked onto fresh BHI agar plates and incubated for 24 hours at 37 °C. Colony PCR was then used to amplify a 1,400 bp region of the bacterial 16S rRNA gene using the universal primers 27F and 1492R [22]. Excess primers and unincorporated nucleotides were removed through a standard ExoSAP-IT reaction. Amplicons were sequenced via Sanger sequencing using the 1492R primer through Functional Biosciences (Madison, WI). Additional sequence reactions using the 27F primer were performed on select isolates when a full length 16S rRNA sequence was needed for genus level identification. Sequences were trimmed for quality and taxonomic identify was determined via comparison against the Ribosomal Database Project and the NCBI nr database via BLASTN [23,24]. Sequences generated as part of this study are publicly available in the NCBI under accession numbers PQ031467-PQ031946, PQ031296-PQ031342, and PQ031373-PQ031381.

### Enrichment of Salmonella and E. coli O157

For specific enrichment of *Salmonella*, aliquots of internal and external fly-bacterial suspensions were diluted two-fold in 30% glycerol and streaked onto both Xylose Lysine Tergitol-4 (XLT4) and MacConkey agar plates. An additional 300 μl of each suspension was also inoculated into 3 ml of iodine-activated tetrathionate (TT) broth, incubated for 24 hours at 37°C, and streaked onto XLT4 and MacConkey agar [25,26]; 100ul of TT cultures were further sub-cultured in 10 ml Rappaport-Vassiliadis (RV) broth for 24 hours at 37 °C and streaked onto XLT4 and MacConkey agar. Following a 24-hour incubation at 37 °C, plates from all three enrichment methods were visually inspected for the presence of black colonies on XLT4 or colorless colonies (lactose non-fermenting) on MacConkey agar. Colonies were tested for both hydrogen sulfide production and the lack of lactose fermentation by heating colony suspensions at 65°C for 15 minutes. Phenotypic identification was validated via PCR to amplify a 284 bp segment of the *invA* gene with *Salmonella* specific primers [27]. Typing of confirmed *Salmonella* isolates was determined using an antisera agglutination test against C1 (O:7) and K (O:18) serogroups and performed according to manufacturer’s instructions (Cedarlane Labs, Burlington, NC).

Selective enrichment for O157 serogroup *E. coli* was performed via immunomagnetic separation using Dynabeads anti-*E. coli* O157 (Applied Biosystems, Waltham, MA) in accordance with standard manufacturer’s protocols. Briefly, 350 μl of each fly-bacterial suspension was diluted tenfold into tryptic soy broth and incubated for 18 h with shaking (200 rpm) at 37°C. A 1 ml aliquot of overnight enrichment culture was mixed with 20 μl of magnetic anti-*E. coli* O157 beads. Suspensions were loaded into a magnetic plate, and non-attached cells were removed from the suspension through triplicate washes (1x PBS- + 0.05% Tween 20). Magnetic beads were resuspended in 100 μl of wash buffer, serially diluted in 1x PBS, and plated onto MacConkey agar with Sorbitol, Cefixime and Tellurite (CT-SMAC) [28]. Following an overnight incubation at 37°C, colorless colonies were isolated and identified through Sanger sequencing of 16S rRNA gene as described above.

### Statistical analyses

Statistical differences in bacterial taxa incidence rates between trap location (barn or calf hutch area), sample type (internal or external), or time (early or late season) were determined using Fisher’s exact test implemented in R (version 4.2.1) using the rstatix package [29,30].

## Results

### Fly Sample pools and homogenate plating

A total of 23 fly pools (230 flies total) consisting of whole *Stomoxys* flies retrieved from adhesive fiberglass traps placed at the Emmons Blaine Dairy Cattle Center in Arlington, WI, were processed to generate both internal and external associated bacterial homogenates. Sixteen pools were composed of flies collected from traps set around the perimeter of two free-stall barn structures; the remaining seven pools were composed of flies collected from a single trap near a calf hutching area.

### Incidence rates of clinically relevant bacterial taxa

Fly homogenates were plated on nutrient agar, and bacterial growth was observed for all samples. A total of 537 bacterial colonies, comprised of 303 internally derived isolates and 234 externally derived isolates, were analyzed via Sanger sequencing of the 16S rRNA gene. We identified 315 sequences aligned to Gram-negative bacteria, of which 54 were determined to be from taxa within the order Pseudomonadales and 239 from the order Enterobacterales (Table 1), which included several genera within the family *Enterobacteriaceae*: *Enterobacter*, *Cronobacter*, *Citrobacter*, *Kosakonia, Leclercia, Escherichia*, *Salmonella*, and *Klebsiella*. Less-frequently isolated taxa included members of the Burkholderiales, Hyphomicrobiales, and Xanthomonadales. Notably, the four Hyphomicrobiales sequences, identified from two separate internal fly pools, most closely aligned with strains of *Brucella-Ochrobactrum*. We additionally identified 222 sequences aligned to Gram-positive bacteria, of which 202 were determined to be from taxa with the order Bacillales (Table 2). A small number of bacterial colonies were identified as belonging to the order Lactobacillales, which included strains of *Aerococcus*, *Desemzia*, *Enterococcus*, and *Lactococcus*. Enrichment plating for *Salmonella* yielded a single isolate from an internal fly pool. A positive agglutination reaction against an O:18 antigen confirmed that the tested isolate belonged to serogroup K. *E. coli* O157 was not detected in any sample using immunomagnetic separation enrichment techniques.

**Table 1.** Taxonomic assignment and number of different gram-negative bacterial colonies isolated from fly sample pools.

| Order | Family | Genus | Isolates |
| --- | --- | --- | --- |
| Burkholderiales | *Alcaligenaceae* | *Alcaligenes* | 9 |
|  | *Comamonadaceae* | *Delftia* | 1 |
| Enterobacterales | *Enterobacteriaceae* | *Cedecea* | 1 |
|  |  | *Citrobacter* | 3 |
|  |  | *Enterobacter* | 46 |
|  |  | *Escherichia* | 47 |
|  |  | *E. coli* O157<sup>1</sup> | 0 |
|  |  | *Klebsiella* | 20 |
|  |  | *Kosakonia* | 44 |
|  |  | *Leclercia* | 4 |
|  |  | *Pseudescherichia* | 1 |
|  |  | *Salmonella* O:18<sup>2</sup> | 1 |
|  |  | *Siccibacter* | 1 |
|  | *Erwiniaceae* | *Erwinia* | 8 |
|  |  | *Pantoea* | 56 |
|  | *Yersiniaceae* | *Serratia* | 8 |
| Hyphomicrobiales | *Brucellaceae* | *Brucella-Ochrobactrum* | 4 |
| Pseudomonadales | *Pseudomonadaceae* | *Pseudomonas* | 31 |
|  | *Moraxellaceae* | *Acinetobacter* | 23 |
| Xanthomonadales | *Xanthomonadaceae* | *Stenotrophomonas* | 8 |

<sup>1</sup>*Escherichia* O157 isolation was performed through immunomagnetic separation of overnight enrichment cultures
<sup>2</sup>Potential isolates were confirmed via amplification of *invA* using *Salmonella* specific primers and serogroup was determined via an antiserum agglutination test.

**Table 2.** Taxonomic assignment and number of different gram-positive bacterial colonies isolated from fly sample pools.

| Order | Family | Genus | Isolates |
| --- | --- | --- | --- |
| Bacillales | *Bacillaceae* | *Cytobacillus* | 4 |
|  |  | *Alkalihalobacillus* | 6 |
|  |  | *Bacillus* | 69 |
|  |  | *Exiguobacterium* | 3 |
|  |  | *Lysinibacillus* | 2 |
|  |  | *Oceanobacillus* | 5 |
|  |  | *Paenibacillus* | 2 |
|  |  | *Peribacillus* | 3 |
|  | *Planococcaceae* | *Bhargavaea* | 1 |
|  | *Staphylococcaceae* | *Mammaliicoccus* | 30 |
|  |  | *Staphylococcus* | 77 |
| Lactobacillales | *Aerococcaceae* | *Aerococcus* | 1 |
|  | *Carnobacteriaceae* | *Desemzia* | 1 |
|  | *Enterococcaceae* | *Enterococcus* | 8 |
|  | *Streptococcaceae* | *Lactococcus* | 2 |
| Micrococcales | *Dermabacteraceae* | *Brachybacterium* | 1 |
|  | *Micrococcaceae* | *Glutamicibacter* | 2 |
|  |  | *Kocuria* | 2 |
|  |  | *Leucobacter* | 1 |
|  |  | *Microbacterium* | 1 |
|  | *Promicromonosporaceae* | *Cellulosimicrobium* | 1 |

The distribution of select taxa identified across sample pools is shown in Tables 3-5. Among Gram-negative bacteria, we infrequently detected *Acinetobacter*, *Serratia*, and *Enterococcus* across both barn and calf area fly samples. Conversely, *Pantoea*, *Pseudomonas*, and taxa within the *Enterobacteriaceae* were widespread among sample pools. Notably, *Escherichia* strains were detected in 21.7% of internal fly pools and 26.1% of external fly pools; *Klebsiella* was detected in 13.0% of internal fly pools and 17.4% of external fly pools (Table 6). No significant differences in incidence rates were found between sample pool types, trap location, or sampling time for either *Escherichia* or *Klebsiella* (*p* > 0.05; Fisher’s exact test). For Gram-positive bacteria, *Bacillus* was detected in 82.6% of internal pools and 69.6% of external pools; taxa within the *Staphylococcaceae*, including *Staphylococcus* and *Mammaliicoccus,* were also commonly isolated from all samples. However, while no significant differences in incidence rates between sample pool types, trap location, or sampling time were observed for *Bacillus* (Table 7) (*p* > 0.05; Fisher’s exact test), barn samples harbored a significantly higher incidence of *Staphylococcaceae* than calf samples (*p* = 0.02; Fisher’s exact test). The incidence of *Staphylococcaceae* was also significantly higher in internal barn samples than in external barn samples (*p* = 0.04; Fisher’s exact test) (Table 7), although the same statistical trends did not apply for tests conducted with incidence rates for *Staphylococcus* or *Mammaliicoccus* at the genus level.

**Table 3.** Distribution of select clinically relevant bacterial taxa cultured from flies captured early season near free-stall barns.

|  |  | Internal Pools |  |  |  |  |  |  |  | External Pools |  |  |  |  |  |  |  |
| --- | --- | --- | --- | --- | --- | --- | --- | --- | --- | --- | --- | --- | --- | --- | --- | --- | --- |
| Taxa |  | 1 | 2 | 3 | 4 | 5 | 6 | 7 | 8 | 1 | 2 | 3 | 4 | 5 | 6 | 7 | 8 |
| Gram<br>(-) | *Enterobacter* | Y | - | - | Y | Y | - | - | - | Y | - | - | Y | Y | - | - | - |
|  | *Escherichia* | - | Y | - | - | - | - | Y | - | - | Y | - | - | - | - | - | - |
|  | *Klebsiella* | - | - | - | - | - | - | - | - | - | - | - | - | - | - | - | - |
|  | *Kosakonia* | - | Y | Y | - | Y | - | Y | Y | - | - | Y | Y | Y | - | Y | Y |
|  | *Pantoea* | Y | Y | Y | Y | Y | Y | Y | Y | Y | - | Y | Y | Y | - | Y | Y |
|  | *Acinetobacter* | - | Y | - | - | - | - | Y | - | - | - | - | - | Y | - | - | Y |
|  | *Pseudomonas* | Y | Y | Y | Y | Y | - | Y | Y | Y | - | Y | Y | Y | - | - | Y |
| Gram<br>(+) | *Bacillus* | Y | Y | Y | Y | - | Y | Y | - | Y | Y | Y | - | Y | Y | Y | Y |
|  | *Mammaliicoccus* | Y | Y | Y | Y | - | - | Y | Y | - | Y | - | - | Y | - | - | - |
|  | *Staphylococcus* | Y | - | - | - | Y | Y | Y | Y | - | - | - | Y | Y | Y | Y | Y |
|  | *Enterococcus* | - | - | - | - | - | - | - | - | - | - | Y | - | - | Y | Y | - |

**Table 4.**
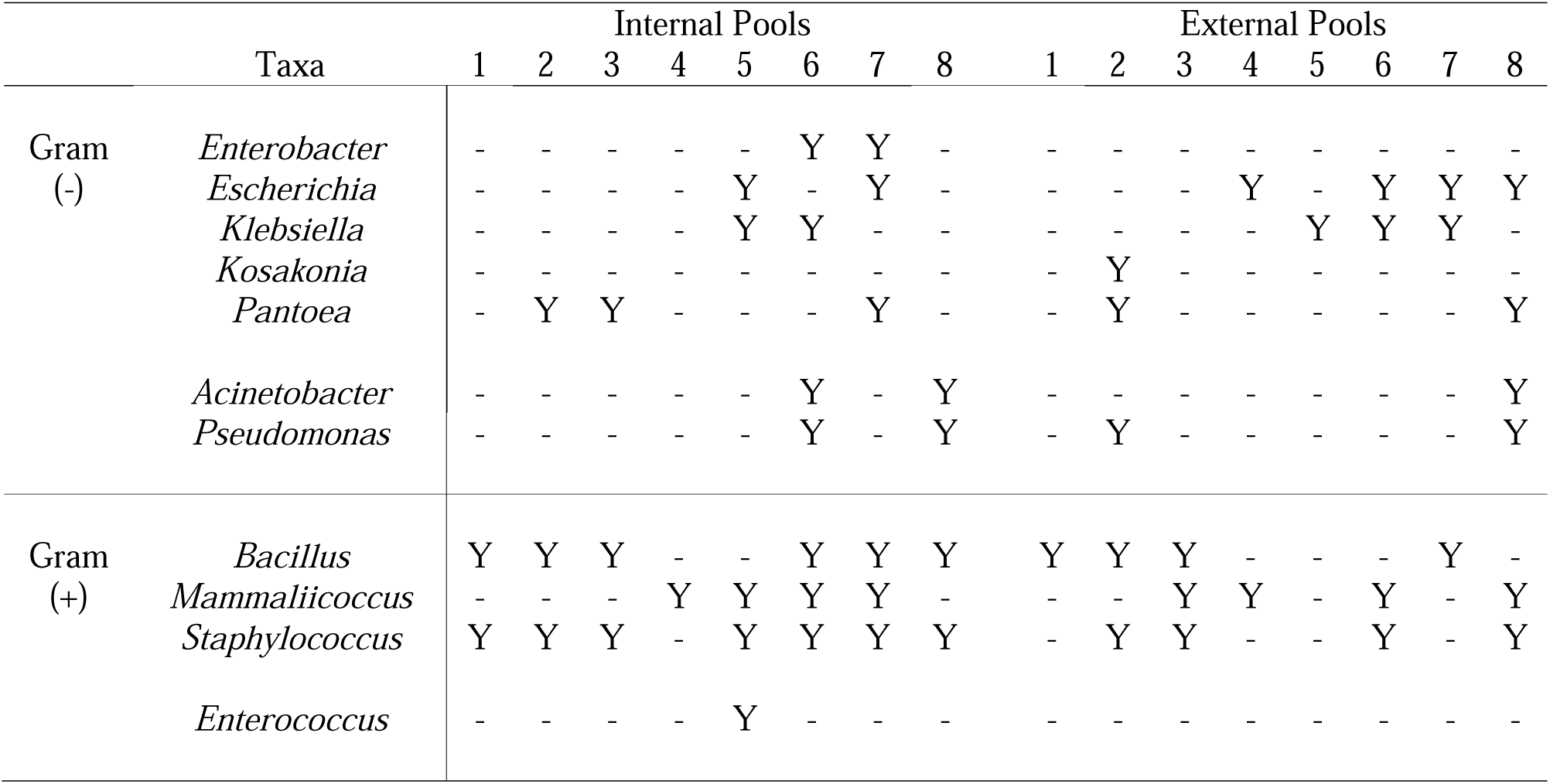
Distribution of select clinically relevant bacterial taxa cultured from flies captured late season near free-stall barns.

**Table 5.**
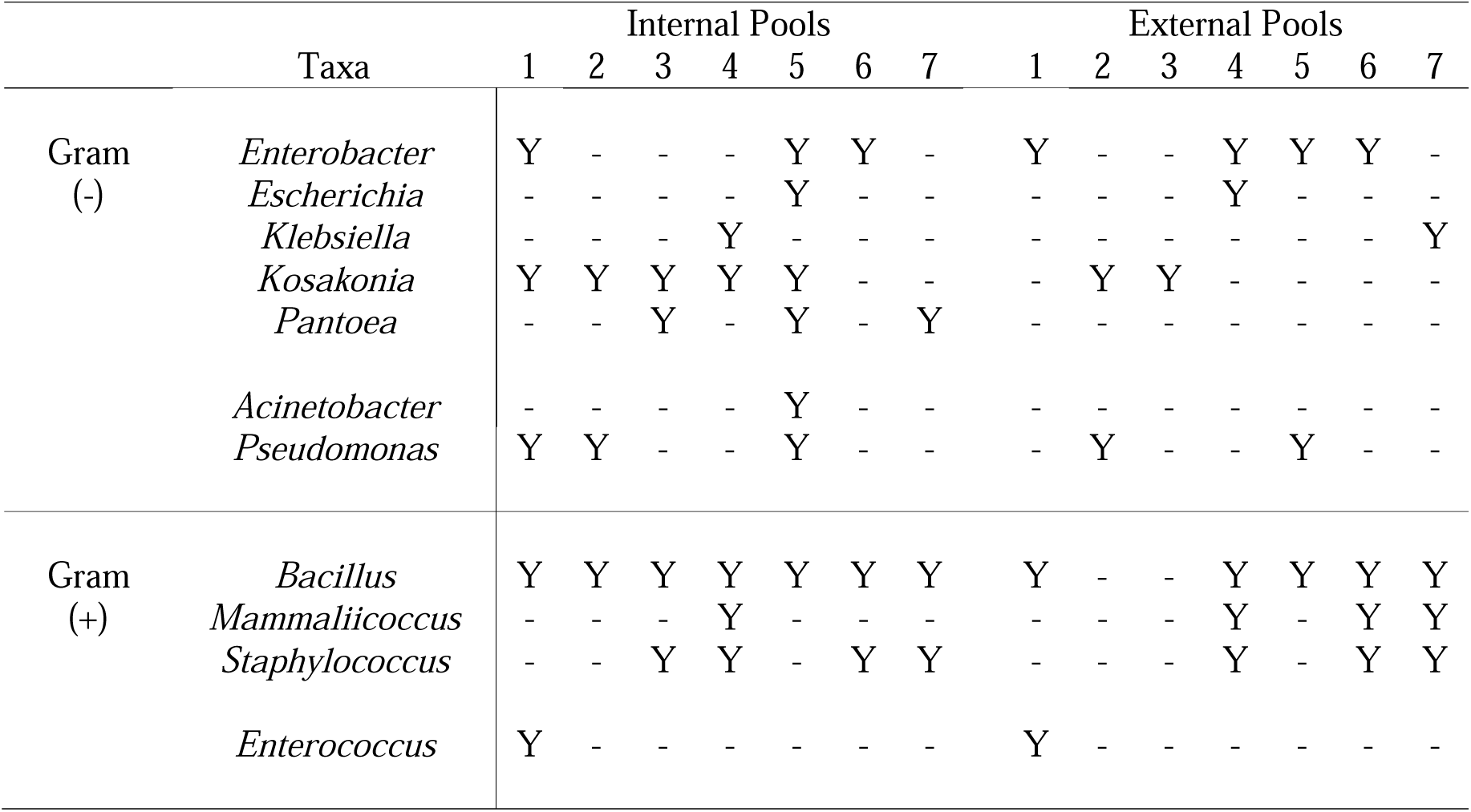
Distribution of select clinically relevant bacterial taxa cultured from flies captured near calf hutches.

|  |  | Internal Pools |  |  |  |  |  |  | External Pools |  |  |  |  |  |  |
| --- | --- | --- | --- | --- | --- | --- | --- | --- | --- | --- | --- | --- | --- | --- | --- |
| Taxa |  | 1 | 2 | 3 | 4 | 5 | 6 | 7 | 1 | 2 | 3 | 4 | 5 | 6 | 7 |
| Gram<br>(-) | *Enterobacter* | Y | - | - | - | Y | Y | - | Y | - | - | Y | Y | Y | - |
|  | *Escherichia* | - | - | - | - | Y | - | - | - | - | - | Y | - | - | - |
|  | *Klebsiella* | - | - | - | Y | - | - | - | - | - | - | - | - | - | Y |
|  | *Kosakonia* | Y | Y | Y | Y | Y | - | - | - | Y | Y | - | - | - | - |
|  | *Pantoea* | - | - | Y | - | Y | - | Y | - | - | - | - | - | - | - |
|  | *Acinetobacter* | - | - | - | - | Y | - | - | - | - | - | - | - | - | - |
|  | *Pseudomonas* | Y | Y | - | - | Y | - | - | - | Y | - | - | Y | - | - |
| Gram<br>(+) | *Bacillus* | Y | Y | Y | Y | Y | Y | Y | Y | - | - | Y | Y | Y | Y |
|  | *Mammaliicoccus* | - | - | - | Y | - | - | - | - | - | - | Y | - | Y | Y |
|  | *Staphylococcus* | - | - | Y | Y | - | Y | Y | - | - | - | Y | - | Y | Y |
|  | *Enterococcus* | Y | - | - | - | - | - | - | Y | - | - | - | - | - | - |

**Table 6.** Proportion of mastitis associated gram-negative bacteria cultured across different sample pool types.

|  | *Enterobacteriaceae* | *Escherichia* | *Klebsiella* | *Pantoea* | *Pseudomonas* |
| --- | --- | --- | --- | --- | --- |
| Barn | 0.813 | 0.438 | 0.188 | 0.750 | 0.625 |
| Calf | 1.000 | 0.286 | 0.286 | 0.429 | 0.429 |
| *p*-value | 0.53 | 0.66 | 0.62 | 0.18 | 0.65 |
| Early (ES) | 0.917 | 0.250 | 0.083 | 0.750 | 0.750 |
| Late (LS) | 0.818 | 0.545 | 0.364 | 0.545 | 0.364 |
| *p*-value | 0.59 | 0.21 | 0.15 | 0.40 | 0.10 |
| Internal | 0.696 | 0.217 | 0.130 | 0.609 | 0.522 |
| External | 0.870 | 0.261 | 0.174 | 0.348 | 0.391 |
| *p*-value | 0.28 | 1.00 | 1.00 | 0.14 | 0.55 |
| Internal Barn | 0.625 | 0.250 | 0.125 | 0.688 | 0.563 |
| External Barn | 0.813 | 0.313 | 0.188 | 0.500 | 0.438 |
| *p*-value | 0.43 | 1.00 | 1.00 | 0.47 | 0.72 |
| Internal Calf Area | 0.857 | 0.143 | 0.143 | 0.429 | 0.429 |
| External Calf Area | 1.000 | 0.143 | 0.143 | 0.000 | 0.286 |
| *p*-value | 1.00 | 1.00 | 1.00 | 0.19 | 1.00 |
| Internal-ES | 0.917 | 0.167 | 0.083 | 0.750 | 0.750 |
| Internal-LS | 0.455 | 0.273 | 0.182 | 0.455 | 0.273 |
| *p*-value | 0.03* | 0.64 | 0.59 | 0.21 | 0.04* |
| External-ES | 0.917 | 0.167 | 0.000 | 0.500 | 0.500 |
| External-LS | 0.818 | 0.273 | 0.364 | 0.182 | 0.273 |
| *p*-value | 0.59 | 0.64 | 0.04* | 0.19 | 0.40 |
| Internal-ES Barn | 0.875 | 0.250 | 0.000 | 1.000 | 0.875 |
| Internal-LS Barn | 0.375 | 0.250 | 0.250 | 0.375 | 0.250 |
| *p*-value | 0.12 | 1.00 | 0.47 | 0.03* | 0.04* |
| External-ES Barn | 0.875 | 0.125 | 0.000 | 0.750 | 0.625 |
| External-LS Barn | 0.750 | 0.500 | 0.375 | 0.250 | 0.250 |
| *p*-value | 1.00 | 0.28 | 0.20 | 0.13 | 0.31 |

Table shows the proportion of a given bacterial taxon cultured in different fly pool groups alongside the corresponding *p*-values for each comparison. Fly pools were grouped by timeframe of capture (early versus late season), location of capture (barn versus the calf hutch area), and fly sample type (isolated from internal versus external fly surfaces). \**p*-value < 0.05; Fisher's exact test.

**Table 7.** Proportion of mastitis associated gram-positive bacteria cultured across different sample pool types.

|  | *Bacillus* | *Enterococcus* | *Mammaliicoccus* | *Staphylococcus* | *Staphylococcaceae* |
| --- | --- | --- | --- | --- | --- |
| Barn | 0.875 | 0.250 | 0.813 | 0.813 | 1.000 |
| Calf | 1.000 | 0.143 | 0.429 | 0.571 | 0.571 |
| *p*-value | 1.00 | 1.00 | 0.14 | 0.32 | 0.02* |
| Early (ES) | 1.000 | 0.333 | 0.667 | 0.667 | 0.833 |
| Late (LS) | 0.818 | 0.091 | 0.727 | 0.818 | 0.909 |
| *p*-value | 0.22 | 0.32 | 1.00 | 0.64 | 1.00 |
| Internal | 0.826 | 0.087 | 0.478 | 0.696 | 0.870 |
| External | 0.696 | 0.174 | 0.391 | 0.522 | 0.609 |
| *p*-value | 0.49 | 0.67 | 0.77 | 0.37 | 0.09 |
| Internal Barn | 0.750 | 0.063 | 0.625 | 0.750 | 1.000 |
| External Barn | 0.688 | 0.188 | 0.375 | 0.563 | 0.688 |
| *p*-value | 1.00 | 0.60 | 0.29 | 0.46 | 0.04* |
| Internal Calf Area | 1.000 | 0.143 | 0.143 | 0.571 | 0.571 |
| External Calf Area | 0.714 | 0.143 | 0.429 | 0.429 | 0.429 |
| *p*-value | 0.46 | 1.00 | 0.56 | 1.00 | 1.00 |
| Internal-ES | 0.833 | 0.083 | 0.583 | 0.583 | 0.833 |
| Internal-LS | 0.818 | 0.091 | 0.364 | 0.818 | 0.909 |
| *p*-value | 1.00 | 1.00 | 0.41 | 0.37 | 1.00 |
| External-ES | 0.750 | 0.333 | 0.250 | 0.500 | 0.583 |
| External-LS | 0.727 | 0.000 | 0.455 | 0.545 | 0.636 |
| *p*-value | 1.00 | 0.09 | 0.40 | 1.00 | 1.00 |
| Internal-ES Barn | 0.750 | 0.000 | 0.750 | 0.625 | 1.000 |
| Internal-LS Barn | 0.750 | 0.125 | 0.500 | 0.875 | 1.000 |
| *p*-value | 1.00 | 1.00 | 0.61 | 0.57 | 1.00 |
| External-ES Barn | 0.875 | 0.375 | 0.250 | 0.625 | 0.750 |
| External-LS Barn | 0.500 | 0.000 | 0.500 | 0.500 | 0.625 |
| *p*-value | 0.28 | 0.20 | 0.61 | 1.00 | 1.00 |

Table shows the proportion of a given bacterial taxon cultured in different fly pool groups alongside the corresponding *p*-values for each comparison. Fly pools were grouped by timeframe of capture (early versus late season), location of capture (barn versus the calf hutch area), and fly sample type (isolated from internal versus external fly surfaces). \**p*-value < 0.05; Fisher's exact test.

## Discussion

*Stomoxys* and other synanthropic flies, which develop in bovine manure and other decaying organic materials, are suspected mechanical vectors of microbial pathogens [2,3,31]. While much attention is given to the microbiota of houseflies (*M. domestica*), particularly for its role in the transmission of bacterial pathogens [7,32], far less is known about the native microbiota of *Stomoxys* flies, which come in frequent contact with dairy cattle or other mammalian hosts during required nutritional bloodmeals. In this study, we present the first comprehensive survey showing the frequency and distribution of culturable bacteria associated with *Stomoxys* flies. Our findings indicate that *Stomoxys* flies harbor culturable bacteria, including *Enterobacteriaceae*, *Pseudomonas*, *Bacillus*, and *Staphylococcus* spp. that can be readily isolated from both internal and external surfaces. Notably, the most frequently identified bacterial taxa are commonly associated with diseases in both humans and animals, including bovine mastitis. The high prevalence of such clinically relevant bacteria, paired with universally large fly populations on dairy barns, suggests a possible role for *Stomoxys* flies in the transmission or dispersal of environmental pathogens.

A major goal of this study was to determine if detected culturable bacteria differed between internal and external fly samples, trap location, or time of capture. As the dipteran microbiota harbors a low-complexity microbiota dominated by a few highly-abundant taxa [33], culturing methodologies can used to reasonably estimate carriage rates of fly-associated bacterial taxa. However, it is likely that our results may be an underrepresentation of the true incidence rates, and our methodology does not account for any obligate anaerobes, obligate intracellular pathogens, or other bacteria that are unculturable on standard microbiological growth media. Nevertheless, we found, with only a few exceptions, no statistically significant differences in incidence rates of commonly occurring bacterial taxa between trap location, sample type, or time. These results align with our previous 16S rRNA amplicon sequencing data on the fly microbiota, indicating that the fly-associated microbial communities are primarily influenced by fly life history and physiology [18].

We note that this study focused on fly populations on a single dairy farm during peak fly season, and it is unknown if the fly microbiota is influenced by larger seasonal changes in weather conditions, which influences farming practices and bovine behavior. Flies are also highly mobile, with the potential to move among or between farms [6]. Potential differences between calf and barn collected flies may therefore have been minimized due to the proximity of the barn to the calf hutching area, where fly populations may simultaneously interact with both adult cows and calves.

Of particular interest is the potential role of flies in the spread of bovine mastitis, a bacterial infection of the udder tissues characterized by the inflammation of the mammary glands. Host immune responses, including recruitment and proliferation of leukocytes to the affected mammary gland, reduce milk quality in lactating cows [34]. Except for host-adapted strains of *Staphylococcus aureus* and *Streptococcus agalactiae*, mastitis pathogens are primarily environmentally derived, with opportunistic infections occurring after exposure of the cow teat to manure or soiled bedding [12]. Non-contagious strains of *Staphylococcus* and *Streptococcus* are the most frequently isolated Gram-positive environmental mastitis pathogens [12]. Within our samples, we identified a high proportion of fly pools with culturable *Staphylococcaceae* bacteria with sequences matching to NAS belonging to *S. saprophyticus/xylosus* and *Mammaliicoccus sciuri* (previously classified as *S. sciuri*) clusters. While NAS are often identified as one of the most frequent causes of persistent subclinical mastitis with the potential to induce clinical mastitis symptoms, there remains little research on the exact contribution of NAS to udder health and the pathogenicity of different NAS taxa is not well understood [35–37]. We additionally detected a high frequency of *Bacillus* across sample pools, which are known minor mastitis pathogens occasionally isolated from milk [38]. While many *Bacillus* are considered nonpathogenic or commensals in humans, strains of *Bacillus cereus* are significant human pathogens, causing both gastrointestinal infections (*B. cereus*) and anthrax (*B. anthracis*) [39]. In contrast to *Staphylococcus* and *Bacillus*, we did not detect culturable *Streptococcus* in any of our fly samples and other Lactobacillales (*Enterococcus*, *Lactococcus*, and *Aerococcus*) were identified only infrequently. Our previous 16S rRNA amplicon sequencing study also found *Streptococcus* to be rare in *Stomoxys* flies, suggesting that the *Stomoxys* fly gut may not be suitable for the growth of *Streptococcus*. Culture-independent studies, including our previous work, have conversely identified reads assigned to *Enterococcus* bacteria as prevalent across internal *Stomoxys* fly microbiota [18,40]. Interestingly, other culture-based studies have simultaneously observed low incidence rates of *Enterococcus* bacteria in *Stomoxys* flies [16,17]. This discrepancy can be partly explained by a reduction of bacterial viability during sample collection and subsequent freezing [41,42]. Alternatively, reads assigned as *Enterococcus* from culture-independent studies could represent DNA sequenced from dead or nonviable cells [43]. Low incidences of culturable *Enterococcus* could therefore also be driven by selective pressures, including the enzymatic lysis of cells within the fly gut [44].

Among Gram-negative bacteria, *Enterobacteriaceae* are most frequently identified as causative agents of environmentally-acquired bovine mastitis [12]. We identified several culturable *Enterobacteriaceae* common to fly pools, which included *Enterobacter*, *Kosakonia*, *Escherichia*, and *Klebsiella.* Both *Klebsiella* and *E. coli* are important mastitis pathogens, and often associated with acute clinical cases [45]. *E. coli* isolates derived from mastitic milk are genomically diverse and share no distinct evolutionary clade [46,47]. Instead, host factors are regarded as the primary drivers of infection outcomes [48]. Less is known about the pathogenesis of *Klebsiella* and other *Enterobacteriaceae*, although structural lipopolysaccharides expressed on the cell surface of Gram-negatives can trigger an inflammatory immune response in host udder tissues [49]. This suggests that fly-derived *Enterobacteriaceae* have the capacity to cause oppurtinistic mastitis infections; however, *in vivo* studies would be required to confirm the pathogenic potential of isolates.

Across samples, we detected other Gram-negative taxa, including *Acinetobacter*, *Pseudomonas*, *Serratia*, and *Pantoea*, which are known to be associated with both bovine mastitis and opportunistic infections in humans [38,49,50]. *Brucella* was also detected across two internal fly pools, which showed the strongest sequence similarity to reference strains previously classified as *Ochrobactrum* [51]. While highly pathogenic lineages such as *B. abortus* are a major causative agent of abortion and metritis in lactating dairy cattle [52], strains previously classified as *Ochrobactrum* represent free-living opportunistic pathogens [51]. Further work will be needed to understand if *Stomoxys* also participate in the carriage of obligate pathogenic *Brucella* strains. In contrast to mastitis-associated bacterial taxa, we identified low incidence rates of the enteric pathogens Salmonella and E. coli O157, despite the use of specific enrichment protocols. We note that fly samples used in this study were collected from a prior field collection study and stored at −20 °C for a period of approximately 2 years. It is possible that cold storage of samples and freeze-thaw cycles resulted in a decrease in the number of viable culturable colonies, as has been previously reported [53]. This suggests that the observed incidence rates, especially for taxa with an initially low abundance, could be an underestimate. Alternatively, low incidence rates of enteric pathogens could be a result of fly life-history. Blood-feeding by stable flies, as has been previously shown in mosquitos, likely acts as a strong selective pressure on the gut microbiome, reducing overall microbial diversity [54]. Coprophagous muscid flies, which come into frequent contact with manure and continually re-uptake manure-associated bacteria, may show higher incidence rates of certain manure borne pathogens. Past research on beef cattle farms have observed greater carriage rates of E. coli O157 in house (M. domestica), face (M. autumnalis), and blow (Family: Calliphoridae) flies, relative to stable flies [55,56]. While Salmonella in flies has been previously reported [57–59], prevalence rates often vary greatly between studies. These differences highlight the need for further comparative studies of how fly life history and environmental conditions impact their potential role in the carriage and potential dissemination of bacterial pathogens.

## Conclusions

In this study, we determined the incidence rates of culturable microbes from the internal and external surfaces of biting flies. We found that flies harbor a high abundance of culturable *Staphylococcus* and Enterobacterales, suggesting the potential for transmission of pathogenic microbes by flies on a dairy barn environment. Our results highlight the need for further research to understand how management practices, life history, and geographic conditions impact the fly microbiota. The combined usage of culture-dependent methodologies and sequence-based technologies will also be necessary to determine the functional capacity and genetic diversity of pathogens carried by *Stomoxys* flies. Such studies could have significant implications for public health and infectious disease control on dairy farms.

## List of abbreviations

ETEC: enterotoxigenic Escherichia. Coli
EHEC: enterohemorrhagic E. coli
NAS: non-aureus staphylococci
PBS: phosphate-buffered saline
EMB: Eosin-Methylene Blue
BHI: Brain Heart Infusion
PCR: polymerase chain reaction
XLT4: Xylose Lysine Tergitol-4
TT: iodine-activated tetrathionate
RV: Rappaport-Vassiliadis
CT-SMAC: MacConkey agar with Sorbitol, Cefixime and Tellurite

## Declarations

### Ethics approval and consent to participate

Not applicable

### Consent for publication

Not applicable

### Availability of data and materials

Data supporting the conclusions of this article, along with scripts used for analysis and table generation, are available in the Coon laboratory GitHub repository [60]. Sequences generated as part of this study are publicly available in the NCBI GenBank (https://www.ncbi.nlm.nih.gov/genbank/) under accession numbers PQ031467-PQ031946, PQ031296-PQ031342, and PQ031373-PQ031381.

### Authors’ contributions

Sample collection was performed by AJS and KLC. All authors contributed to the design of the methodology; AJS, CDL, and ADJT processed samples for bacterial culturing and identification. AJS wrote the initial draft, and KLC, GS, CDL, and ADJT contributed to revisions.

## Acknowledgements

We thank Jessica Cederquist and the UW-Madison Department of Animal and Dairy Science Dairy Herd Operation for access to the Emmons Blaine Arlington Dairy Research Center.

